# Rapid termite diversification is associated with increased transposable element activity

**DOI:** 10.64898/2026.02.08.704128

**Authors:** Cong Liu, Simon Hellemans, Alina A. Mikhailova, Cédric Aumont, Yi-Ming Weng, Aleš Buček, Jan Šobotník, Mark C. Harrison, Dino P. McMahon, Thomas Bourguignon

## Abstract

Termites are a lineage of social insects that originated during the Early Cretaceous ∼150 million years ago. They are the dominant decomposers in modern tropical and subtropical terrestrial ecosystems, a role they achieved through several phases of rapid diversification facilitated by unknown genetic mechanisms. Here, we investigated the link between termite diversification and the activity of transposable elements (TEs). We reconstructed the evolutionary history of TE replications using the genomes of 45 termite species and identified two waves of TE expansion that involved all major TE classes and superfamilies and took place synchronously across termite lineages. The first wave occurred around the end of the Cretaceous, and the second wave occurred during the Oligocene and Miocene, coinciding with the two major phases of termite diversification. We further estimated TE insertion/deletion rates along the species tree and showed that TE activity is positively correlated with termite diversification during the last ∼150 million years of evolution, providing evidence for a link between TE activity and diversification over a macroevolutionary timescale.

## Introduction

Termites, the second largest lineage of social insects, are represented by ∼3,000 species primarily distributed across tropical and subtropical terrestrial ecosystems (Krishna et al. 2013; Jones and Eggleton 2011). They account for 10 to 20% of the animal biomass in tropical ecosystems (Ellwood and Foster 2004; Eggleton et al. 1996; Rosenberg et al., 2023), where they are the dominant macroscopic decomposers of organic matter (Griffiths et al. 2019), from sound wood to humus and soil (Donovan et al. 2001; Bourguignon et al. 2011). Termites were coined by Wilson (1987) as “the little things that run the world” for their role as ecosystem engineers, having major effects on soil properties and plant growth in modern terrestrial ecosystems (Jouquet et al. 2011, 2016; Evans et al. 2011; Wu et al. 2025).

The timing of the rise of termites is increasingly well understood. The origin of termites is estimated at ∼150 million years ago (mya), around the Jurassic and Cretaceous boundary (Bourguignon et al. 2014; Bucek et al. 2019). The fossil record shows that termites already formed a diverse group ∼100 mya, during the mid-Cretaceous (Thorne et al. 2000; Engel et al. 2009, 2016; Zhao et al. 2021). However, Neoisoptera and Kalotermitidae, two sister lineages that diverged ∼120 mya (Bourguignon et al. 2014; Bucek et al. 2019) and make up 99% of modern termite species (Krishna et al. 2013), are poorly represented among mid-Cretaceous termite fossils. Modern termite diversity originated more recently in two phases of rapid diversification (Hellemans et al. 2025). The first phase involved Kalotermitidae and Neoisoptera around the end of the Cretaceous (∼65 mya), while the second phase involved multiple lineages of Neoisoptera, predominantly Termitidae, after the Eocene-Oligocene boundary (∼34 mya) (Heimburger et al. 2022; Hellemans et al. 2025). The second phase of rapid termite diversification coincides with their ecological diversification, including the evolution of new feeding niches, particularly soil-feeding in Termitidae (Hellemans et al. 2022a), and the acquisition of a global distribution (Bourguignon et al. 2016, 2017; Bucek et al. 2022). Termites also experienced morphological diversification during this period, especially the Termitidae, which evolved new soldier defensive strategies (Prestwich 1984) and complex worker hindgut structures (Noirot 2001). The genetic mechanisms behind termite diversification are unknown.

One category of genetic elements associated with rapid species diversification is transposable elements (TEs) (Lönnig et al. 2002). TEs are a major component of eukaryotic genomes (Feschotte and Pritham 2007; Elliott and Gregory 2015; Osmanski et al. 2023). They were originally viewed as selfish elements spreading within genomes at the expense of their host’s fitness (Doolittle and Sapienza 1980; Orgel and Crick 1980). However, TE activity, which may increase under environmental shifts and some stress factors, generates genetic variation that can act as a substrate for evolutionary forces, potentially facilitating evolutionary innovations (Belyayev 2014; Dubin et al. 2018; Schrader and Schmitz 2019; Naciri and Linder 2020). For example, TE activity has been linked to the diversification of various organisms, such as mammals (Ricci et al. 2018), fishes (Brawand et al. 2014; Auvinet et al. 2018), and *Drosophila* (Craddock 2016; Oliveira et al. 2024). TE activity has also been shown to facilitate adaptation to novel environments in the peppered moth (Hof et al. 2016) and invasive ants (Schrader et al. 2014; Errbii et al. 2021). While termite TEs have previously been studied in the context of TE defense and longevity (Elsner et al. 2018; Berger et al. 2022; Post et al. 2023; Qiu et al. 2023), their potential role in termite diversification has remained unexplored.

Here, we reconstructed TE activity during termite evolution and demonstrated their link with termite diversification. First, we built histograms of the evolutionary distance among TEs using the genomes of 45 taxonomically diverse termite species and two cockroach outgroups (Liu et al. 2025a). By estimating the mutation rate from fourfold degenerate (4dtv) sites of single-copy orthologous genes, we converted evolutionary distance into calendar time and show that the timing of bursts in TE activity coincides with termite diversification history. Second, we showed that the rates of TE insertion/deletion directly inferred on the branches of a termite phylogenetic tree based on 473,032 TE insertion/deletion characters correlate with previous estimates of net termite species diversification rates (Hellemans et al. 2025).

## Results and discussion

### Most termites experienced two waves of TE bursts

We reconstructed the history of TE expansions for each species individually using histograms representing the evolutionary distance distribution of all TE lineages inferred from sequence alignments (Figures 1A-C, S1). Assuming a constant rate of net TE accumulation, the distribution is expected to be exponential, with most TE lineages being young. TE bursts over short time periods create deviation in the exponential distribution, taking the shape of peaks and platforms. We first attempted to detect peaks and platforms by fitting mixture distributions containing one to ten lognormal components. However, the best models, identified using the Bayesian information criterion, contained 10 components in 34 of the 47 species and always contained at least five components (Figures 1A-C, S1), indicating overfitting, a well-documented phenomenon in mixture model fitting (Naik et al. 2007; Vekemans et al. 2012; Johnson et al. 2016; Tiley et al. 2018). As an alternative approach, we used the significant zero-crossing test of the first derivative of the distributions (Chaudhuri and Marron 1999; Barker et al. 2008; Vanneste et al. 2015), which provided more reliable identification of peaks and platforms (Figures 1A-C, S1). The method identified 77 peaks and platforms amongst the 47 species, each representing a TE burst in the evolutionary history of the corresponding species. The distributions contained a single burst in 18 species, two bursts in 28 species, and three bursts in *Dolichorhinotermes longilabius*.

**Figure 1.**
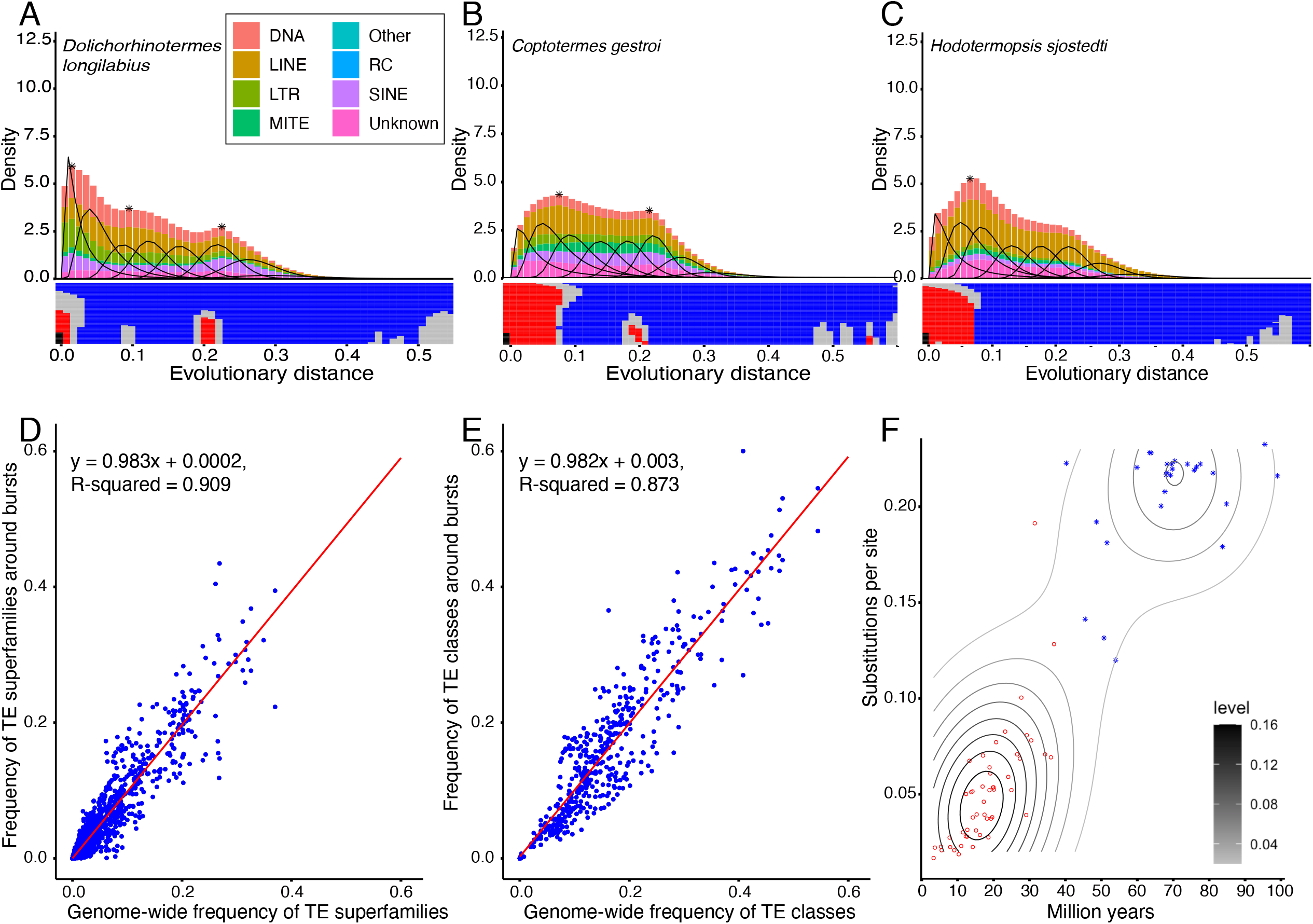
TE bursts found in this study. Histograms representing the probability density distribution of the averaged evolutionary distance between TE lineages for three representative species showing (A) three significant bursts, (B) two significant bursts, and (C) one significant burst. Histograms of all 47 species studied here are available in Figure S1. Bar colors indicate the main TE classes. The black curves inside the histogram show the lognormal mixture model with the best Bayesian Information Criteria values. The stars indicate significant peaks or platforms identified by the significant zero-crossings of the first derivative of the distribution from SiZer. The SiZer maps are represented below the histograms. The rows of SiZer maps represent various bandwidth of the significant zero-crossing testing. The color of tiles shows the monotonicity of the distribution: red for increasing, blue for decreasing, and grey for platform. Regressions between the proportion of each (D) TE superfamily and each (E) TE class around the observed TE bursts and across the whole distribution. (F) Scatter plot showing the evolutionary distance against estimated calendar time of TE bursts, with black contour lines indicating density. The red points represent TE burst events classified as “recent” and the blue points TE burst classified as “ancient.”

TE bursts can result from the increased activity of a few TEs, while most other elements remain silent (Belyayev 2014). To determine whether the TE bursts identified in this study involved a few TEs or were global, involving whole TE landscapes, we performed ordinary least squares (OLS) between the frequency of TE superfamilies/classes at 0.02 substitutions per site around all identified TE bursts and their frequencies along the 47 whole distributions. We detected significant positive correlations in all identified TE bursts for both TE superfamilies and classes (*p* < 10^-15^), with slopes of 0.983 and 0.982, intercepts that are not significantly different from 0 (*p* > 0.15), and R-squared of 0.909 and 0.873, indicating that the frequency of each TE category around bursts closely resemble their overall frequency (Figures 1D-E). Furthermore, when adding TE superfamily and class as a second variable in the OLS, no significant effect was detected by an analysis of variance (ANOVA) (*p* > 0.676). These results indicate that TE bursts were not driven by extensive replications of specific superfamilies or classes but were global across termite TE landscapes.

### Two waves of TE bursts took place independently in most termite lineages

We estimated the age of TE bursts to determine whether they occurred in parallel across many termite species or were ancient, occurring in the ancestor of modern termites and leaving a signature in the genomes of its descendants. Assuming TE sequences accumulate substitutions neutrally (Barrón et al. 2014; Arkhipova 2018), the evolutionary distance between two TE lineages can be translated into absolute time by using the neutral substitution rate, which is equivalent to the single-nucleotide mutation rate (Kimura et al. 1971). We estimated the substitution rate of fourfold degenerate (4dtv) sites of 1,410 single-copy orthologous genes along the entire species tree and used it as a proxy for the mutation rates of the two cockroach and 45 termite genomes analyzed here. The substitution rates for termite and *Cryptocercus* terminal branches ranged from 6.9 x 10^-10^ to 3.02 x 10^-9^ substitutions per site per year and were 1.93 x 10^-9^ substitutions per site per year for *B. orientalis* (Figure S2). These estimations are generally similar to vertebrates (Kumar and Subramanian 2002; Zhang et al. 2020; Bergeron et al. 2023) and an order of magnitude lower than estimates for other insects (Peccoud et al. 2017). This likely reflects the long generation time of termites (Li and Wu 1987), which is comparable to vertebrates and much longer than most insects (Tasaki et al. 2021). These estimated mutation rate values across the termite tree of life allow us to date the waves of TE bursts.

Two clusters of TE bursts appeared when plotting the substitutions per site against the estimated calendar time in million years of the 77 peaks and platforms identified by the significant zero-crossing tests (Figures 1F, 2A). These clusters correspond to two waves of TE bursts that we delimit based on calendar time estimations but slightly overlap in terms of substitutions per site. The evolutionary distances of the more recent wave vary between 0.02 and 0.19 substitutions per site (median: 0.050), corresponding to between 3.3 and 36.8 mya (median: 17.4). The evolutionary distances of the more ancient wave vary between 0.12 and 0.23 substitutions per site (median: 0.22), which corresponds to 40.2 and 99.0 mya (median: 68.3). The recent wave of TE bursts was observed in all investigated species except *Mastotermes darwiniensis*, and the ancient wave was detected in 26 of 47 species (Figure S1). In some of the remaining 21 species, such as *Hodotermopsis sjostedti, Stolotermes victoriensis, Microcerotermes* sp., *Silvestritermes heyeri*, and *Cylindrotermes parvignathus* (Figure S1), platforms matching the ancient wave of TE bursts could be visually identified at around 0.2-0.22 substitutions per site (54.7-98.8 mya), suggesting that the ancient wave of TE bursts also influenced most termite lineages, but its signal fell below the detection threshold. The presence of two waves of TE bursts across termites indicates dramatic changes in TE activity along geological time.

The two waves of TE bursts took place in parallel across termite lineages (Figure 2A). Among the 50 TE bursts making up the recent wave, only seven were placed on internal branches of the time-calibrated species tree, while others were placed on terminal branches, indicating independent events across the species examined here. The ancient wave of TE bursts occurred independently in the ancestors of several termite lineages, although their precise locations on the tree are more difficult to ascertain. Seven internal branches and five terminal branches were associated with the ancient wave of TE bursts, representing multiple bouts of TE expansion across several lineages, including Mastotermitidae, Archotermopsidae, Kalotermitidae, and Neoisoptera, although some of these 12 branches may represent the same events. For example, the ancient TE bursts that were placed on the branches leading to Termitidae, Heterotermitidae + Termitidae (Geoisoptera), Psammotermitidae, and Geoisoptera + Psammotermitidae likely represent fewer than four events and may even represent a single event that was imprecisely located across multiple adjacent branches in the tree. Overall, our results demonstrate that at least two waves of TE bursts occurred in most termite lineages in parallel.

**Figure 2.**
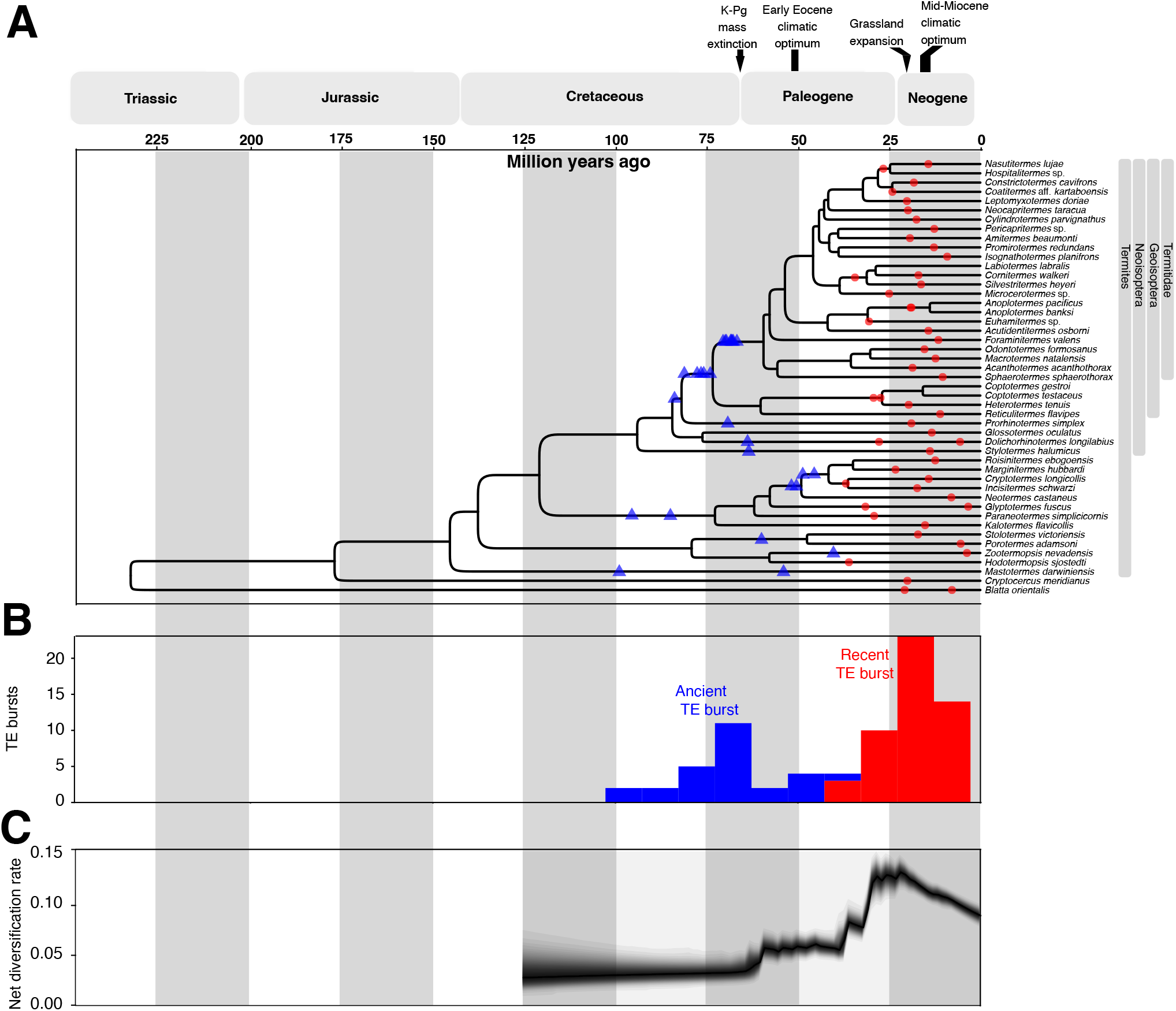
(A) Time-calibrated species tree of termites showing the location of TE bursts. Red circles and blue triangles indicate the recent and ancient waves of TE bursts, respectively. (B) The distribution of TE bursts along time. (C) The net diversification rate of termites against time, adapted from Hellemans et al. (2025).

### The two waves of TE bursts coincided with periods of rapid termite diversification during major environmental changes

The recent wave of TE bursts took place during the Oligocene and Miocene (33.9-5.3 mya) (Figure 2). This period was characterized by a global cooling of the Earth’s climate (Zachos et al. 2001; Goldner et al. 2014), the retraction of megathermal rainforests towards the tropics and subtropics (Morley 2011), and the expansion of grasslands (Retallack 2001; Edwards et al. 2010; Berggren and Prothero 2014). Concurrently, termites rapidly diversified during the Oligocene and Miocene, giving rise to most modern termite genera (Heimburger et al. 2022; Hellemans et al. 2025). Termites, especially Neoisoptera, also colonized the world through several dozen transoceanic dispersal events (Lee et al. 2015; Bourguignon et al. 2016, 2017; Hellemans et al. 2022a) and became the dominant decomposers in the tropics during this period (Engel et al. 2009).

The timing of the ancient wave of TE bursts spread over a 60-million-year period, with the mode of the distribution occurring ∼70 mya, at the end of the Cretaceous (Figure 2). The origin of termites dates back to ∼150 mya (Engel 2007; Bucek et al. 2019), predating the ancient wave of TE bursts. Termites were a diversified insect group during the Cretaceous; however, they were primarily represented by extinct families or extant but species-depauperate families during that period (Thorne et al. 2000; Engel et al. 2009, 2016; Zhao et al. 2021; Barden and Engel 2021). The ancestors of extant termites experienced a phase of rapid diversification at the end of the Cretaceous (Hellemans et al. 2025), which coincided with the ancient wave of TE bursts inferred herein from the genomes of extant termites. Therefore, akin to the recent wave, our results suggest that the ancient wave of TE activity coincided with a period of increased termite diversification following the emergence of new environments after the Cretaceous mass extinction. Altogether, our results suggest a link between termite diversification and TE activity.

### Net termite diversification rates positively correlate with TE activity

So far, we showed that TE bursts coincided with spikes of termite diversification (Figure 2); however, we did not formally test for a correlation. Next, we tested for a correlation between TE insertion/deletion rates and termite net diversification rates calculated for all branches of four time-calibrated phylogenetic trees inferred from 2,800 termite samples, differing in the combination of nuclear and mitochondrial sequences used for their reconstruction (Hellemans et al. 2025). The trees were reduced to 45 operational taxonomic units (OTUs) representing termite species included in this study (Table S1). The topologies of the four trees were almost identical to those of Liu et al. (2025a) (Figure 2A). Net diversification rates were inferred for each tree by Hellemans et al. (2025) using BAMM (Rabosky et al. 2014a). The rates of TE insertion/deletion were the ratios between the branch lengths of trees inferred from the presence/absence of 473,032 TEs located near orthologous ultraconserved elements and the branch lengths of the four corresponding time-calibrated trees. Spearman’s correlation test showed significant correlation (*p* < 0.003) between net species diversification rate and the rates of TE insertion/deletion in all four trees, with coefficients between 0.33 and 0.50. This correlation was also supported by OLS without branches shorter than one million years, for which the estimations of TE insertion/deletion rate can be noisy (all *p* < 0.0001), and up to 34.8% variation of net species diversification rate was predictable from TE insertion/deletion rate (Figure 3). Therefore, species diversification is linked to TE activity over geological time.

**Figure 3.**
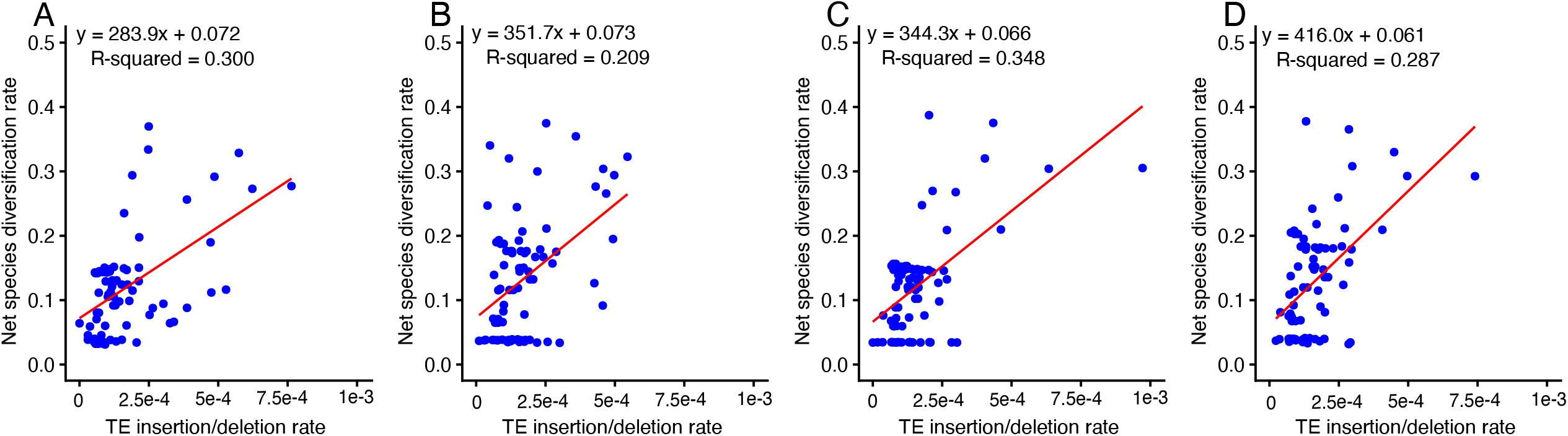
Regression plots showing the relationship between TE insertion/deletion rates and net species diversification rates for species tree branches longer than one million years. The rates were estimated using trees reconstructed in Hellemans et al. (2025), with (A) mitochondrial genomes with third codon positions included, (B) mitochondrial genomes without third codon positions, (C) UCEs and mitochondrial genomes with third codon positions included, and (D) UCEs and mitochondrial genomes without third codon positions.

TE activity affects genome evolution in multiple ways, ranging from local changes of gene structures and regulatory elements to large-scale genomic rearrangements, all of which have complex effects on phenotypes and provide the raw material on which natural selection can operate (Casacuberta and Gonzalez, 2013; Lisch 2013; Belyayev 2014; Arkhipova 2018; Schrader and Schmitz 2019; Naciri and Linder 2020). While the adaptive role of TEs has been reported in some species (Schrader et al. 2014; Errbii et al. 2021; Hof et al. 2016; Esnault et al. 2019; Rech et al. 2022; Ren et al. 2025), our understanding of the role of TE activity at macroevolutionary time scales is limited to some reports that TE bursts coincide with evolutionary radiations (Pace and Feschotte 2007; Brawand et al. 2014; de Boer et al. 2007) and correlation between TE activity and extant lineage diversity (Ricci et al. 2018). Here, we went further and explicitly inferred rates of TE insertion/deletion and species diversification on termite phylogenetic trees, and showed that TE activity is positively correlated with termite diversification over the past 150 million years.

### Conclusions

We identified two waves of parallel TE bursts in many termite lineages that involved all major TE classes and superfamilies. These two waves of TE bursts coincided with two bursts of termite diversification at the end of the Cretaceous and during the Oligocene-Miocene period. We developed a new approach to measure TE insertion/deletion rates on all branches of a phylogenetic tree and demonstrated that it positively correlates with net species diversification rates in termites. Our results suggested that TEs have contributed to termite diversification on a macroevolutionary timescale (∼150 million years).

## Materials and Methods

### Genome assemblies and annotations of TEs

We used the genome assemblies of 45 termites and two cockroaches produced by Liu et al. (2025a). We used the TE annotation of Liu et al. (2025b), which classified TEs populating each genome into 37,966 families belonging to all major classes, such as miniature inverted-repeat TEs (MITEs), DNA transposons, long terminal repeats (LTRs), long interspersed nuclear elements (LINEs), and short interspersed nuclear elements (SINEs) (Figure S3). All TEs were retained, including fragmented TEs. Annotated repeats that were not TEs were removed.

### Reconstruction of the history of TE replication in termite genomes

We reconstructed the evolutionary history of TE replications for each species separately. We first clustered TE fragments belonging to each TE family using CD-HIT v4.8.1 (Li and Godzik 2006), which was run with a DNA sequence identity threshold set to 80%. MITEs shorter than 50 bp were not considered for downstream analyses. For other TE classes, sequences shorter than 100 bp were filtered out. The sequences of each cluster containing fewer than 10,000 fragments were aligned using MAFFT v.7.508 (Katoh et al. 2002) with the “*–auto*” option. We did not consider clusters composed of more than 10,000 fragments because of computational limitations. Multiple sequence alignments were used to compute pairwise distances using the *dist*.*dna* function implemented in the R package *ape* (Paradis and Schliep 2019). The *dist*.*dna* function was run using the options “*model=K80, pairwise*.*deletion=TRUE*” to calculate Kimura’s 2-parameter distances (Kimura 1980) and remove sites with missing data in the pairwise comparisons. We used the matrix of pairwise Kimura’s 2-parameter distances to conduct hierarchical clustering with the *hclust* function implemented in the R package *fastcluster* (Müllner 2013). The hierarchical clustering was performed with the option *“method=average,”* producing ultrametric trees from which all node-to-tip distances were extracted to build age distributions of TE replications. Node-to-tip distances represent half the average evolutionary distance — or sequence divergence — between TE sequences descending from the node. We only considered node-to-tip distances varying between 5 x 10^-3^ and 0.5 substitutions per site, that is, nodes with evolutionary distances varying between 0.001 and 1 substitution per site. Since TE sequences are assumed to accumulate point mutations neutrally following their insertions (Lynch and Walsh, 2007), evolutionary distance is a proxy for the age of TE replication (Kimura and Ohta 1971).

We screened the age distributions — or evolutionary distance distributions — of TE replications generated for 45 termites and two cockroaches to identify significant departures from the null model, which assumes a constant rate of TE replication, resulting in an exponential age distribution with most replication events being recent. We expected TE bursts to introduce bumps in the exponential distributions, which can be detected using both parametric and non-parametric methods. We first attempted to model the distribution with mixture models of one to ten lognormal components and fit the age distribution using the expectation-maximization algorithm implemented in the *mixfit* function of the R package *mixR* run with default settings (Yu 2022). Fitted models were evaluated using the Bayesian information criterion. We also used a non-parametric method, the significant zero-crossings (SiZer) (Chaudhuri and Marron 1999) of the first derivative of the kernel density. First, kernel density estimates of the age distribution were computed using the *density* function implemented in R with *bw=“SJ”* (R Core Team, 2022). Then we used R package *SiZer* (Chaudhuri and Marron 1999) to scan the density estimates and produce SiZer map, which shows regions of significant increase or decrease in the distributions (Barker et al. 2008; Vanneste et al. 2015).

Lastly, we investigated whether TE bursts involved specific TEs or the entire TE landscape. We tested the correlation between the representation of each TE superfamily/class along the whole age distribution and within a distance of 0.02 substitutions per site from the peaks and platforms identified with SiZer using the R function *lm*. In order to test if the peaks and platforms were dominant by specific TE superfamilies, we also added TE superfamily/class as a second independent variable in the regression and performed analysis of variance (ANOVA) using the R function *anova*.

### Conversion of evolutionary distances into calendar time

We estimated mutation rates of termites in order to translate evolutionary distances into calendar time (Kimura and Ohta 1971). We first clustered all genes annotated in the 47 genomes by Liu et al. (2025a) into hierarchical orthologous groups (HOGs) using OrthoFinder v.2.5.4 with default settings (Emms and Kelly 2019) and identified 1,410 universally single-copy HOGs. We generated the multiple protein sequence alignment for each single-copy HOG using MAFFT with the option “*–auto*.” Protein alignments were converted into codon alignments using PAL2NAL v.14 (Suyama et al. 2006) and concatenated into a supermatrix from which 41,729 fourfold degenerated (4dtv) sites were extracted. We used the 4dtv sites and the species tree of Liu et al. (2025a) to estimate branch length with IQ-TREE v.2.2.0.3 (Minh et al. 2020). We ran IQ-TREE with the options “*-mset GTR –tree-fix*,” which computed branch length using a GTR+F+R4 model, the best model as determined by ModelFinder (Kalyaanamoorthy et al. 2017). The branch lengths of the 4dtv tree divided by the lengths of corresponding branches in the time-calibrated species tree provide an estimation of mutation rates per year for both internal and terminal branches. We used the mutation rates calculated for all branches of the tree to convert the evolutionary distances of TE replication events into calendar time.

### Estimation of rates of species diversification and TE insertion/deletion

We investigated whether species diversification rates correlate with rates of TE insertion/deletion. Species diversification rates were estimated for four time-calibrated species trees by Hellemans et al. (2025). These four trees contained thousands of tips, including tips corresponding to the termite species included in this study. All following operations were carried out using the R package *BAMMtools* (Rabosky et al. 2014b). We downsampled the tips of these trees and the corresponding species diversification rates to 45 operational taxonomic units (OTUs) corresponding to the termite species included in this study using the *subtreeBAMM* function (Table S1). Net species diversification rates were estimated for each branch as the difference between speciation and extinction rates and extracted using *getMarginalBranchRateMatrix* function. Net diversification curves were obtained using the *plotRateThroughTime* function.

The rate of TE insertion/deletion was estimated using ultraconserved elements (UCEs) identified in Hellemans et al. (2022b) and Liu et al. (2025a). We kept 13,223 UCEs that were located at least 200 bp away from both ends of the contigs in which they were embedded in all 45 termite genomes used in this study. We used presence/absence data of each TE family in the 200-bp flanking every 13,223 UCEs as a proxy for TE insertion/deletion, as described in Liu et al. (2025b). We reasoned that a specific TE family present in the surroundings of a specific UCE generally represents a locus orthologous across termite genomes and can therefore be used as a proxy for TE insertion/deletion. We created a binary matrix using this approach, where each row represented one of the 45 termite species and each column one existing combination of a UCE and a TE family, with presence and absence coded as 0 and 1, respectively. We excluded columns only composed of 0 or 1 (exclusive absence or presence of a TE family near one specific UCE), leaving 473,032 columns. As previously described by Liu et al. (2025b), we confirmed that the binary matrix contained phylogenetic information by inferring a species tree using IQ-TREE with the options “*–seqtype BIN -m MFP -B 1000*.” We used our binary matrix to recompute the branch lengths for the four subsampled timetrees using IQ-TREE with the options “*–seqtype BIN -m MFP –tree-fix*,” which allowed to fix the topology of the trees to that of the time-calibrated species trees of Hellemans et al. (2025). Therefore, recomputed branch lengths reflected the extent of TE insertion/deletion. Finally, we estimated TE insertion/deletion rates along the species tree by dividing the recomputed branch lengths with the corresponding branch lengths of the subsampled time-calibrated trees of Hellemans et al. (2025). Therefore, we obtained TE insertion/deletion rates and net species diversification rates for all branches of the same trees.

### Correlations of the net species diversification rate with rates of TE insertion/deletion

We tested for a correlation between net species diversification rates and the rates of TE insertion/deletion. We repeated the analyses separately for each four species trees built by Hellemans et al. (2025). Spearman’s correlation test was performed using R function *cor*.*test*. We also built linear models using the R function *lm*. Since the estimations of TE insertion/deletion rates for short branches are subject to a high degree of uncertainty, we removed branches shorter than one million years.

## Author contributions

C.L. and T.B. conceptualized the experiments. C.L. and S.H. analysed the data. C.L. and T.B. wrote the manuscript. S.H., A.A.M., C.A., Y.M.W., A.B., J.S., M.C.H., and D.P.M. read and edited the manuscript and accepted the final version.

## Data Availability

The genome assemblies of the 47 species used in this study are available on GeneBank under bioproject PRJNA1198669.

## Acknowledgements

We thank OIST’s Scientific Computation and Data Analysis Section (SCDA) for providing access to the OIST computing cluster.

## Supplementary Material

**Figure S1**. Histogram representing the probability density distribution of the averaged evolutionary distance between TE lineages for all species analysed in this study. Bar colours indicate the main TE classes. The black lines inside the histogram show the lognormal mixture model with the best Bayesian Information Criteria values. The stars indicate significant peaks or platforms identified by the significant zero-crossings of the first derivative of the distribution from SiZer. The SiZer maps are represented below the histograms. The rows of SiZer maps represent various bandwidths of the significant zero-crossing testing. The color of tiles shows the monotonicity of the distribution: red for increasing, blue for decreasing, and grey for platform.

**Figure S2**. Time-calibrated tree of termites showing single-nucleotide mutation rate values estimated for the 47 genomes used in this study. Substitution rates were estimated based on fourfold degenerated (4dtv) sites of 1,410 single-copy orthologous genes. Branch labels represent single-nucleotide mutation rate values per billion years.

**Figure S3**. Repertoires of TE families found across the 47 genomes used in this study. (A) time-calibrated species tree, (B) abundance of TE classes and genome size (star points) in each species, and (C) richness in TE families in each species.

**Table S1**. List of species included in this study and the matching operational taxonomic units and their taxonomic identifications from Hellemans et al. (2025).

