## Supplementary figures and images for "Rapid termite diversification is associated with increased transposable element activity"

### Figure S1

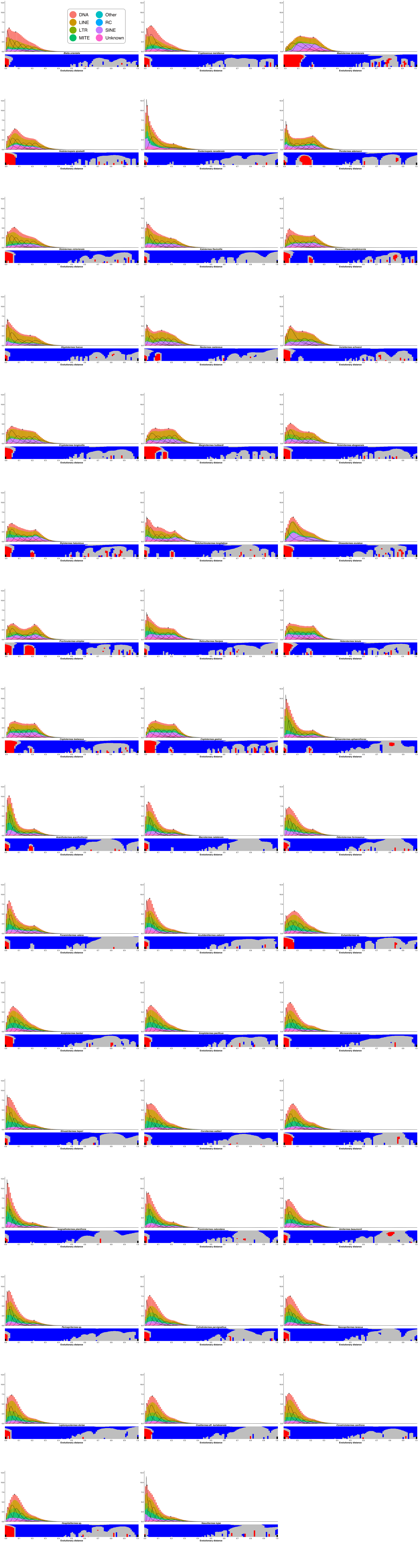

### Figure S2

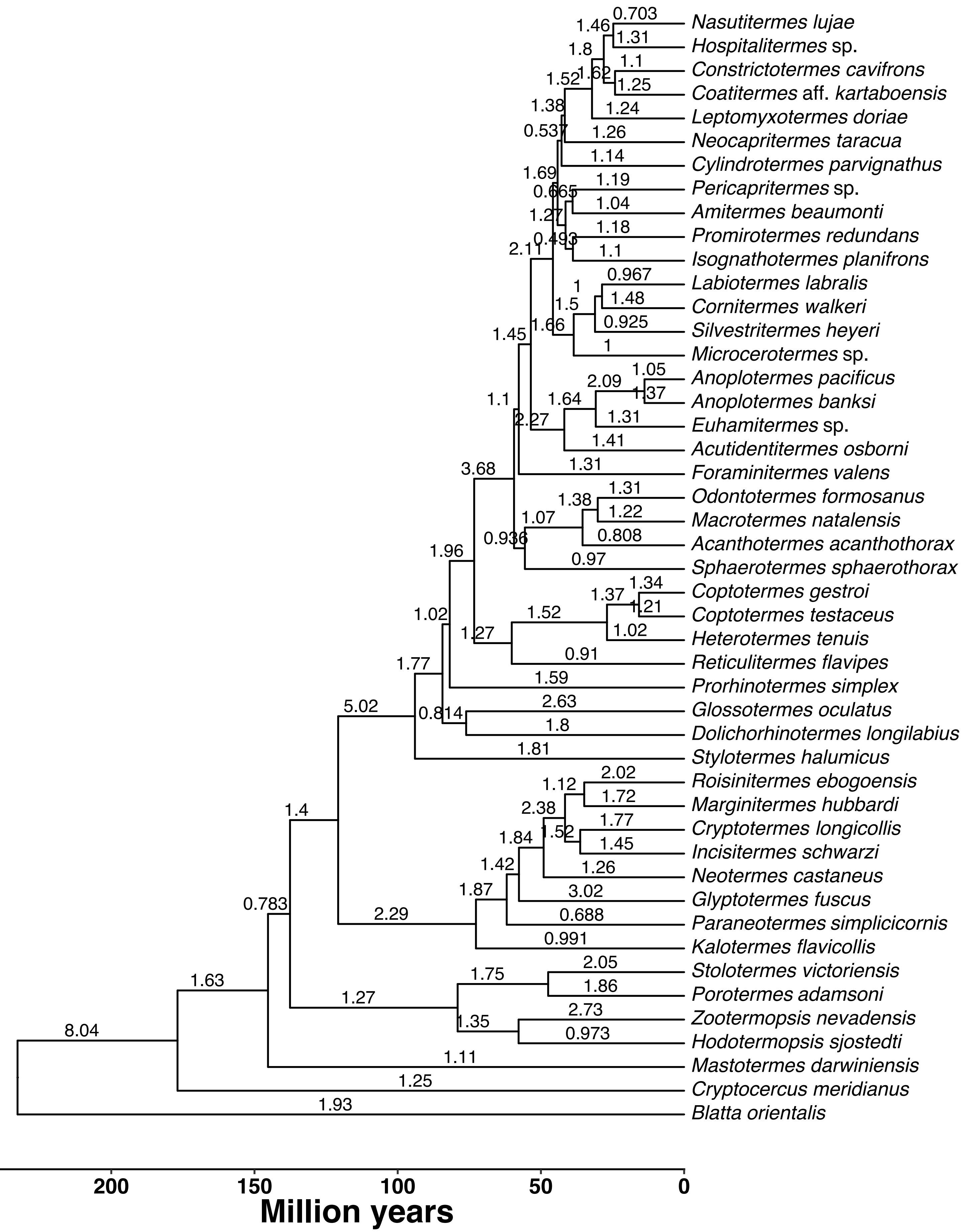

### Figure S3

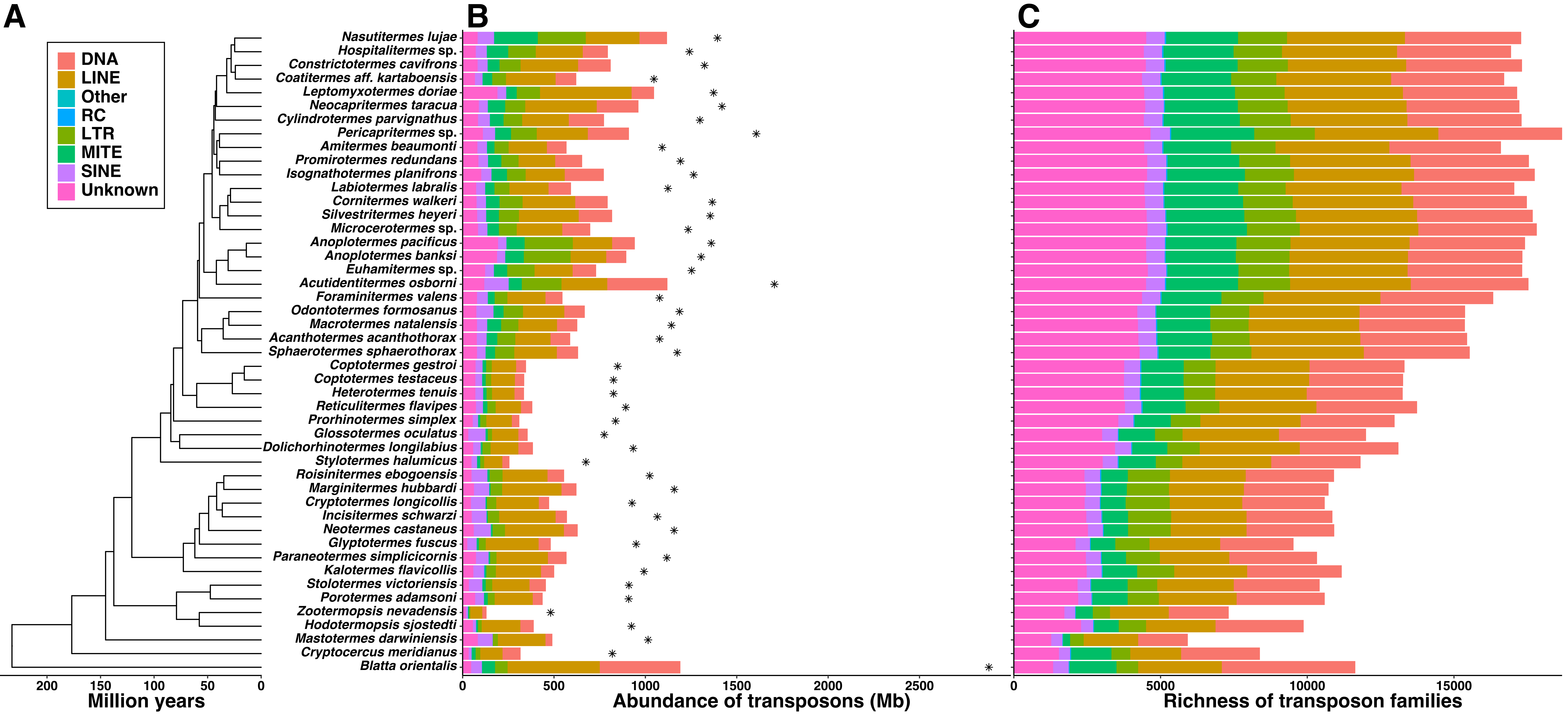
